# Variant calling and genotyping accuracy of ddRAD-seq: comparison with 20X WGS in layers

**DOI:** 10.1101/2024.01.29.577880

**Authors:** Mathilde Doublet, Fabien Degalez, Sandrine Lagarrigue, Laetitia Lagoutte, Elise Gueret, Sophie Allais, Frédéric Lecerf

## Abstract

Whole Genome Sequencing (WGS) remains a costly or unsuitable method for routine genotyping of laying hens methods, thus alternatives have been developed. Among these, reduced representation sequencing approaches can offer both sequencing quality and cost-effectiveness by reducing the genomic regions covered by sequencing. The aim of this study was to evaluate the ability of *double digested Restriction site Associated DNA sequencing* (ddRAD-seq) to identify and genotype SNPs in laying hens, by comparison with a presumed reliable WGS approach. Firstly, the sensitivity and precision of variant calling and the genotyping reliability of ddRADseq were determined. Next, the SNP Call Rate (CR_SNP_) and mean depth of sequencing per SNP (DP_SNP_) were compared between both methods. Finally, the effect of multiple combinations of thresholds for these parameters on genotyping reliability and amount of remaining SNPs in ddRAD-seq was studied. In raw form, the ddRAD-seq identified 349,497 SNPs evenly distributed on the genome with a CR_SNP_ of 0.55, a DP_SNP_ of 11X and a mean genotyping reliability rate per SNP of 80%. Considering genomic regions covered by expected enzymatic fragments (EFs), the sensitivity of the ddRAD-seq was estimated at 32.4% and its precision at 96.4%. The low CR_SNP_ and DP_SNP_ values were explained by the detection of SNPs outside the EFs theoretically generated by the ddRAD-seq protocol. Indeed, SNPs outside the EFs had significantly lower CR_SNP_ (0.25) and DP_SNP_ (1X) values than SNPs within the EFs (0.7 and 17X, resp.). The study demonstrated the relationship between CR_SNP_, DP_SNP_, genotyping reliability and the number of SNPs retained, to provide a decision-support tool for defining filtration thresholds. Severe quality control over ddRAD-seq data allowed to retain a minimum of 40% of the SNPs with a CcR of 98%. Then, ddRAD-seq was defined as a suitable method for variant calling and genotyping in layers.

## Introduction

The development of Next-Generation Sequencing (NGS) approaches has revolutionized genetic marker discovery and genotyping. Depending on the chosen approach, the balance between the marker density, genotype accuracy, the degree of multiplexing of individuals and the experimental costs may vary. Among these approaches, whole genome sequencing (WGS) allows the simultaneous detection and genotyping of the majority of individual polymorphisms at a chosen sequencing depth (1–3). However, it is still challenging mostly because of the sequencing costs per sample. Then, cheaper alternative methods such as low depth WGS have been studied (4,5). Compared to deeper WGS, it offers a large panel of single nucleotide polymorphisms (SNPs) per sample while increasing inter-individual variability and lowering genotyping accuracy (2,6). Moreover, sequencing the whole genome of every individual in a population is often unnecessary, as many biological questions requirering genomic markers (population genetics, genomic selection, genetic diversity studies…) can be answered using only a subset of genomic regions (6).

Then, restriction site associated DNA sequencing (RAD-seq) is a great alternative to WGS. RAD-seq are *de novo* approaches targeting a subset of the genome, thus reducing its complexity and providing a reliable set of markers. These sequencing methods consist in a step of enzymatic digestion followed by different filtration steps applied on the enzymatic fragments (EFs) according to the approach. Then, remaining EFs are amplified by PCR and sequenced to create a library. The diversity of retraction enzymes (REs) available and ways to combine them make RAD-seq methods versatile assay tools (6). Depending on the availability of a good quality reference genome for the species, reads can be aligned thus, improving the proportion of shared markers between individuals (6–8).

The low proportion of shared markers between individuals is a common drawback of RAD-seq approaches. During DNA digestion, RE recognize a specific pattern called restriction sites (RS) to cut DNA. When a mutation or a methylation appears on the DNA, it can either create or delete a RS. Creation of RS for only some of the individuals allows the detection of SNP for only a subset of the studied population thus, introducing missing data between individuals. For creation and deletion of RS, as the mutations occurs only on one strand of the DNA, at an individual level, it means that the two enzymatic fragments generated (one per strand) are not symmetrical on both DNA strands. So, any SNP who would be detected on an unsymmetrical EF would be automatically appear as a homozygote for the SNP allele associated with the successfully digested fragment. Thus, mutations in a RS tends to misestimate genetic diversity within the population (9). PCR bias are also a common bias of RAD-seq approaches leading to a miss estimation of genetic diversity (10). Numerous studies have demonstrated methods to mitigate variant calling and genotyping errors from library preparation to bioinformatics processing of sequencing data (11).

To ensure the lowest possible error rate, all studies systematically apply quality control to variant calling and genotyping data (12). The most common quality control filters are the call rate SNP (CR_SNP_) (13,14) and average sequencing depth per SNP (DP_SNP_) (9,15,16). These filters increase the chances for a SNP to be detected and that allelic frequencies will be well represented in the population. They also ensure to limit the impact of mutations in the RS and methylations on the final data set (9,17). Other filters, such as the probability that the allele frequency distribution respects the Hardy-Weinberg equilibrium (HWE) or the MAF (7) are commonly used to identify and remove SNPs with a high genotyping error rate. The thresholds chosen for each of these filters are rarely justified in the bibliography (18) and depend greatly on the application of the study (6). Some quality control filter thresholds are almost standardized, while others vary from study to study. However, in the case of sequencing methods based on genome reduction, losing genetic information, or wrongly assuming that all SNPs are correct after quality control, can have significant consequences for the conclusions of a study, depending on the application (11,18–22). But, because of the infinite number of events that cause genotyping errors, it is impossible to hope to get rid of them entirely. (23). If not, the studies recommend at least to quantify them in order to give a confidence interval to their results (22–24).

Variant calling and genotyping errors can be quantified through the introduction of replicate sample, by comparing the results of the two sequencings with each other. Variants that are not common to both replicates will be considered erroneous (8,22,24). However, this method cannot identify genotyping errors due to mutations in RS (25) which is a major biases for RAD-seq approaches. Then, the best way to quantify the variant calling and genotyping quality of a sequencing method is to compare it to another more reliable reference method (23,25). The availability of sequencing data reliable enough to be considered as a reference representing the "truth", on the same genomic regions as the RAD-seq method, offers the possibility of calculating the sensitivity and accuracy of the RAD-seq method. Sensitivity corresponds to the ability of the RAD-seq method to detect all the SNPs detected by the reference method, in the genomic regions it covers. Precision, on the other hand, reflects the rate of loci wrongly considered as SNPs by RAD-seq. Sensitivity and accuracy are two indicators of the amount of variant calling and genotyping errors that are rarely found in the literature, and even less so for RAD-seq approaches (3,16) although they have been reported in other studies (26,27). The reliability of sensitivity and precision measurements depends on the quality of the sequencing data used as a reference. In literature, genotypes obtained by a RAD-seq approaches have already been compared to considered “more reliable approaches” such as Sanger sequencing (28), SNP chips (15), or even WGS (25) but never for layers.

Among RAD-seq approaches, double digest Restriction site Associated DNA sequencing (ddRAD-seq) is a method which, thanks to the use of two different RE to digest DNA and a size filtration step for EFs, reduces the inter-individual variability of EFs generated and SNPs detected compared with other RAD-seq methods. It drastically reduces the rate of variant calling and genotyping errors compared with other RAD-seq approaches. The use of two REs also facilitates adapter design and reduces sequencing costs per individual and per base, thus offering the best multiplexing capability compared with other RAD-seq approaches (10,20).Moreover, ddRAD-seq is a sequencing method that can be customized (RE choice, filtering method) to suit the needs of each study (6,20,29). This makes ddRAD-seq a reliable method for variant discovery and genotyping of plants (29–34) and animals (35–39). But, as with other RAD-seq methods, there is no consensus in the literature on the quality control and its filters, according to application, species or protocol features. Furthermore, the thresholds chosen for quality control filters are often not justified or based on other studies. Their real impact on the quality of genotyping data is rarely studied.

But, despite its bias, ddRAD-seq represents a major opportunity for the poultry industry as a small number of markers is sufficient to perform for various applications such as genome-wide association studies (40), linkage disequilibrium calculation (41), CNV detection (42) or genomic selection (43,44) ddRAD-seq is cheaper than deep WGS or HD chips and more accurate than low depth WGS. Compared with LD chips, RAD-seq sequencing enables the integration of the whole genome including micro-chromosomes. These chromosomes are not all represented on commercial chips based on the galGal4 version of the reference genome (Nov, 2011), which integrated only a few of them. The sequencing data can be aligned with the GRCg7b version of the chicken reference genome (January, 2021) in which 39 autosomes and two sexual chromosomes Z and W are represented.

Then, the aim of this study is to assess ddRAD-seq quality of sequencing in terms of variant calling and genotyping, for bi-allelic markers, according to a set of population scale filtering options by comparison with 20X WGS. The 20X WGS was taken as a reference as an effective coverage of 15X is considered as sufficient to achieve high-quality genotyping for WGS(2), The three parts of this work are (i) to describe and compare the SNP calling and genotyping data between WGS and ddRAD-seq, more precisely, (ii) to estimate sensitivity and precision of ddRADseq SNP calling and finally, (iii) to study the genotype concordance of the common SNP between deep WGS and ddRAD-seq.

## Results

### Genome scale parameters

With the 20X WGS, 9,219,123 bi-allelic SNPs were detected on chromosomes 1 to 39 and Z and 51,050 on contigs. With ddRAD-seq, 349,497 bi-allelic SNPs were detected on chromosomes 1 to 39 and Z and 712 on contigs. There were 327,364 SNP common to both methods.

In both cases, at least half of the SNPs (60.9% in 20X WGS and 50.8% in ddRAD-seq) were located on the macro-chromosomes (1-5). Both methods obtained similar results regarding the percentage of SNPs detected on the intermediate chromosomes (6-10) with 15.4% and 16.3% for the 20X and ddRAD-seq respectively. On the contrary, 29.7% of the SNPs identified in ddRAD-seq were located on the micro chromosomes (11-39) against only 19.5% of those found in 20X WGS. Finally, 4.2% of the SNPs from 20X WGS and 3.2% of the SNPs from ddRAD-seq were found on the Z sexual chromosome (Table 1, Fig. 1).

**Fig. 1:**
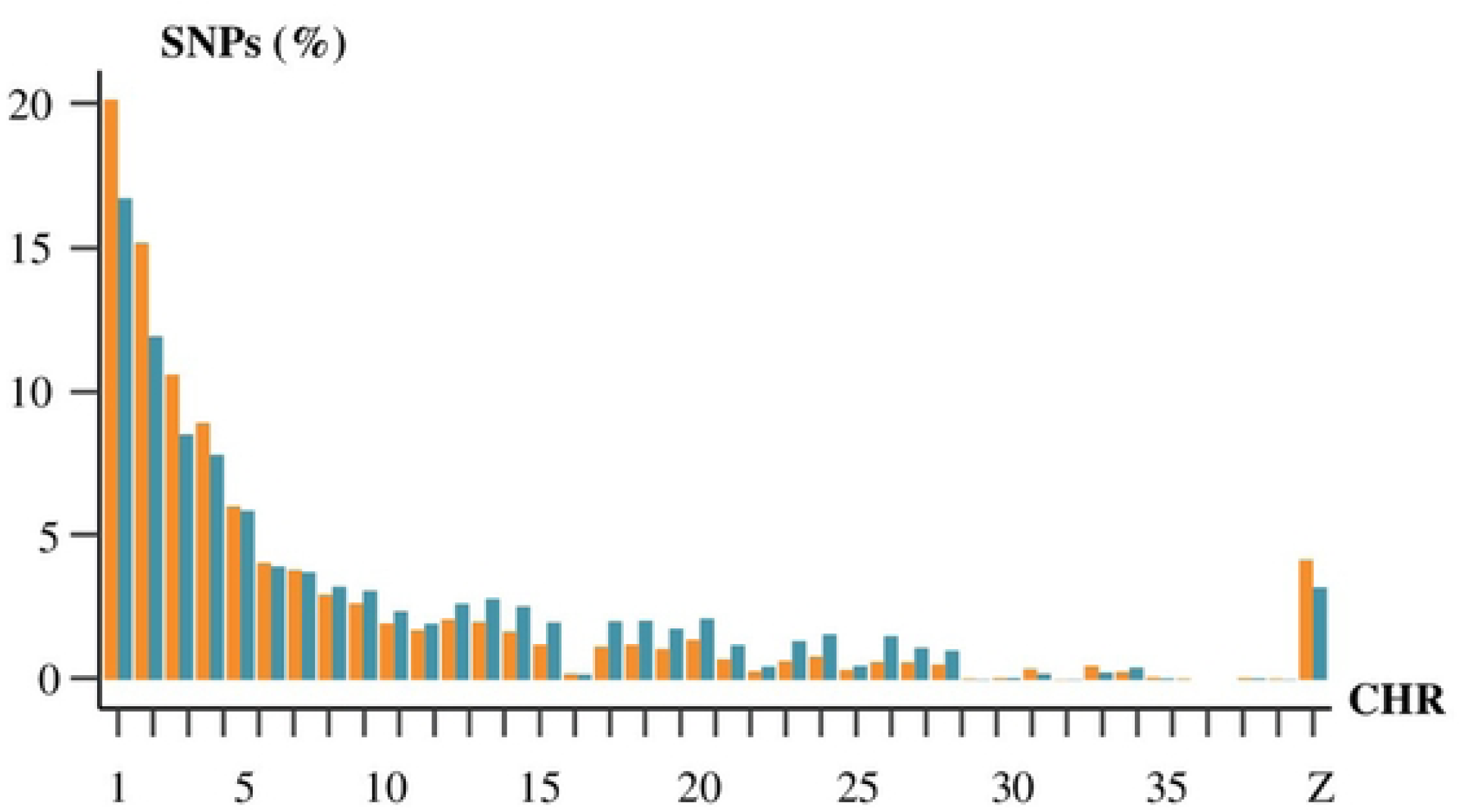
Percentage of total SNP detected by 20X WGS (*in orange*) and ddRAD-seq (*in blue*) on chromosomes 1 to 39 and Z.

**Table 1:** Number of SNPs called in 20X WGS and in ddRAD-seq and their percentage on the total number of SNPs by chromosome type.

|  | *WGS 20X* |  | *ddRAD-seq* |  |
| --- | --- | --- | --- | --- |
|  | #SNPs | %SNPs | #SNPs | %SNPs |
| *Macro (1-5)* | 5 611 994 | 60.9 | 177 658 | 50.8 |
| *Intermediate (6-10)* | 1 420 343 | 15.4 | 56 989 | 16.3 |
| **Micro (11-39)** | 1 803 092 | 19.5 | 103 639 | 29.7 |
| **Z** | 383 694 | 4.2 | 11 211 | 3.2 |

As shown in the Fig 1, the number of SNPs were similarly distributed on chromosomes in 20X WGS or in ddRAD-seq. The number of SNP identified on each chromosome was significantly correlated with the length of each chromosome in 20X WGS (ρ = 0.99) and in ddRAD-seq (ρ = 0.98). The mean distance between two adjacent SNPs called was 3,221 bp and 130 bp for ddRAD-seq and 20X WGS respectively.

### Sensitivity and precision

*In silico* estimation of the genomic regions that should be covered by EFs theoretically generated by the ddRAD-seq protocol was performed as described in the *Materials and Methods*. So, 860,138 Pst1 restriction sites (RS) and 447,997 Taq1 RS were identified on the reference genome GRCg7b. Moreover, 33,600 Pst1 and 63,564 Taq1 RS were created by mutations in our population when considering every individual. A total of 285,004 EFs between 200 and 500 base pair (bp) should have been theoretically generated and pair-end sequenced on an average of 150 bp according to *in silico* prediction.

In 20X WGS, considered here as the reference sequencing approach, 638,135 of the 9,219,123 SNPs were identified inside the expected EFs. Therefore, we expected a sensitivity of 6.9% for ddRAD-seq at a genome-wide scale and a precision surrounding 100%. But, the real sensitivity of ddRAD-seq at a genomic scale was 3.6% and its precision was 93.7%. It represents a loss of sensitivity of 57.9% compared to what was expected.

Then, inside the expected EFs, 214,495 SNPs were identified by ddRAD-seq and 206,872 SNPs were commonly identified by ddRAD-seq and 20X WGS. Using 20X WGS as the reference approach, the effective sensitivity of ddRAD-seq inside the expected EFs was 32.4% and its precision was 96.2%. It represents an even greater loss of sensitivity (67,6%) compared to what was expected then at a genomic scale. This suggested that, in ddRAD-seq, some SNPs were detected outside the expected EFs. Considering the total number of 349,497 SNPs detected in ddRAD-seq, only 61.5% of them were found inside the expected EFs.

### Locations of SNPs outside the EFs

The location of SNPs outside of the expected EFs (*i.e.*, 200 to 500 bp framed by the two enzymatic restriction sites) was investigated as described in Materials and methods. Out of the 134,552 SNPs identified outside the EFs, 18,816 SNPs were found in the 10 bases on each end of these EFs. A total of 47,407 SNPs were located in regions covered by EFs less than 200 bp long but generated by the combination of Taq1 and Pst1. Also, 12 342 SNPs were located in genomic regions covered by Efs cut by Taq1 on both ends and 59 125 by Pst1 on both ends. Additionally, 119,116 SNPs were located in regions corresponding to EFs resulting from the failure of both sizing and selection on RE steps in the ddRAD-seq protocol. Genomic regions concerned by each scenario can overlap with one another. All the effectives of SNPs that could be identified by multiple scenarios overlapping was described in the Fig. 2. Finally, considering each scenario and their overlaps, 4,265 SNPs called out of the EFs were not explained by any of these hypotheses.

**Fig 2:**
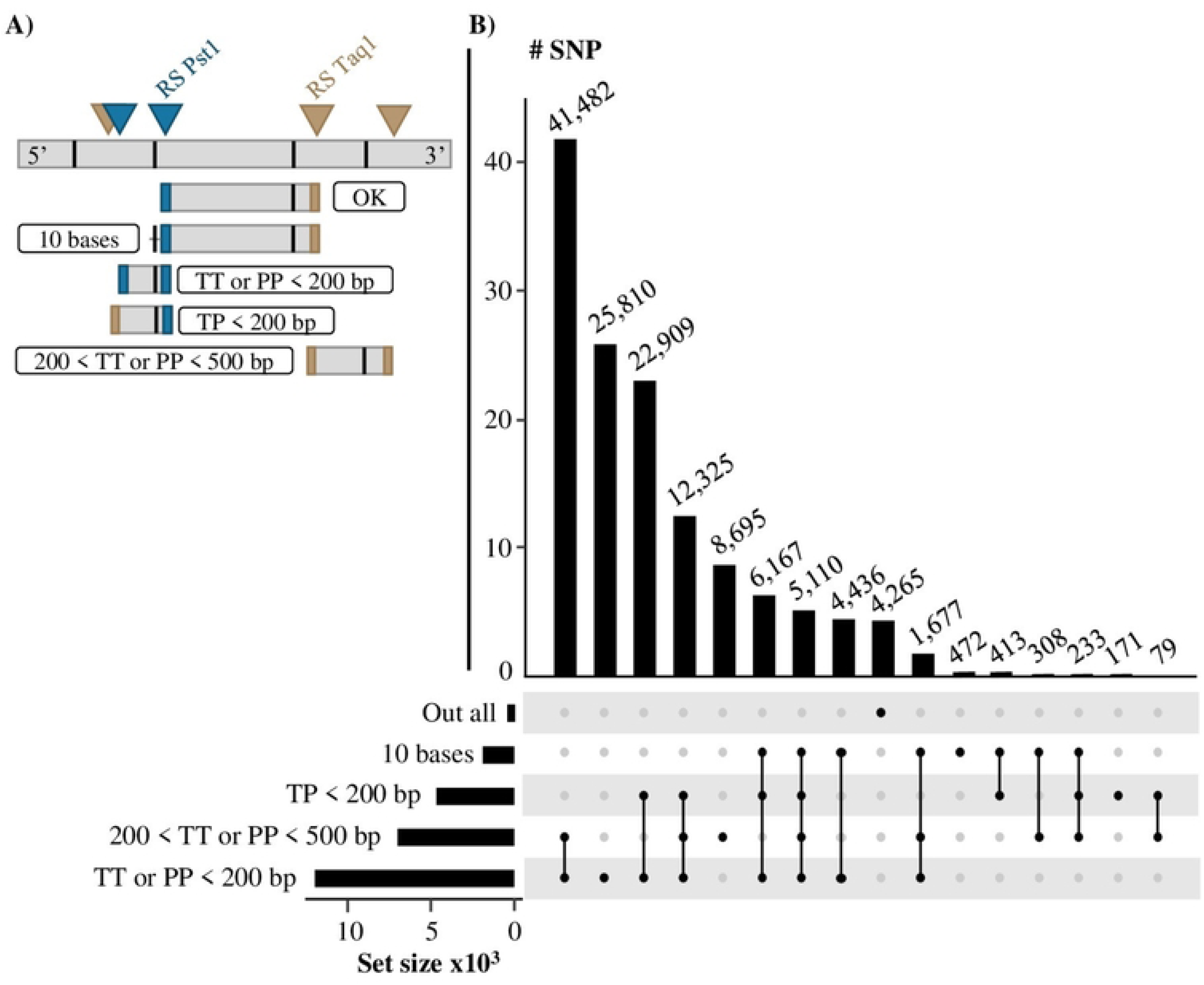
Possible reason for SNPs to be called outside of the theoretical enzymatic fragments theoretically generated by the ddRAD-seq protocol. ***10 bases: SNPs called in the 10 bases on each side out of an enzymatic fragment (EF) between 200 and 500 bp and generated by the combination of Taq1 and Pst1. TP < 200 bp: SNPs located inside the EFs generated by the combination of Taq1 and Pst1 under 200 bp. 200 < TT or PP < 500 bp: SNPs located inside the EFs between 200 and 500 bp, generated by the same restriction enzyme at both ends. TT or PP < 200 bp: SNPs located inside the EFs under 200 bp and generated by the same RS on both sides. Out all: SNPs that don’t fit in any of our scenarios*.**

For ddRAD-seq, the ratio between SNPs inside and outside the theoretical EFs was not different between the chromosomes except for the sexual chromosome Z, where there was as many SNPs inside and SNPs outside the theoretical EFs (Fig. 3). There was no difference of location in specific chromosomic regions between the two sets of SNPs (Supplementary Fig1).

**Fig 3:**
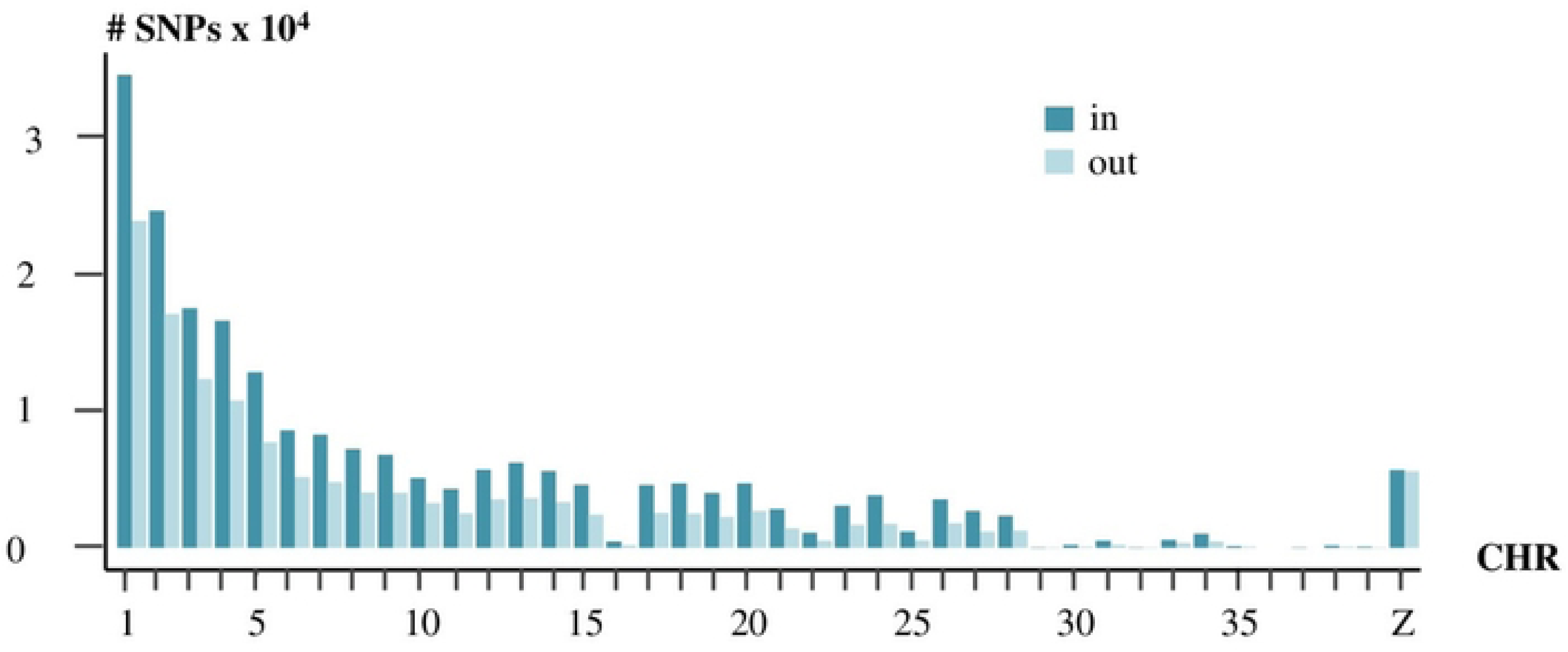
Localization of SNPs in (*in blue*) and out (in light blue) genomic regions theoretically sequenced in ddRAD-seq.

### Effect on CR_SNP_ and DP_SNP_

The CR_SNP_, on average, was significantly higher in 20X WGS (0.99) than in ddRAD-seq (0.55) (p-value < 0,05, Fig. 4A). 95.2% of the SNPs obtained with the 20X WGS were genotyped for all individuals (CR_SNP_=1) while only 29.3% were in ddRAD-seq (Fig. 4B). 56.8% of the ddRAD-seq SNPs had a CR_SNP_ below 0.8 against only 0.9% of the 20X WGS SNPs. Furthermore, the parabolic distribution of CR_SNP_ values in ddRAD-seq showed two higher points, with a large proportion of SNPs having low CR_SNP_ values (0-0.1) and an equivalent proportion having higher CR_SNP_ values (Fig. 4B).

**Fig 4.**
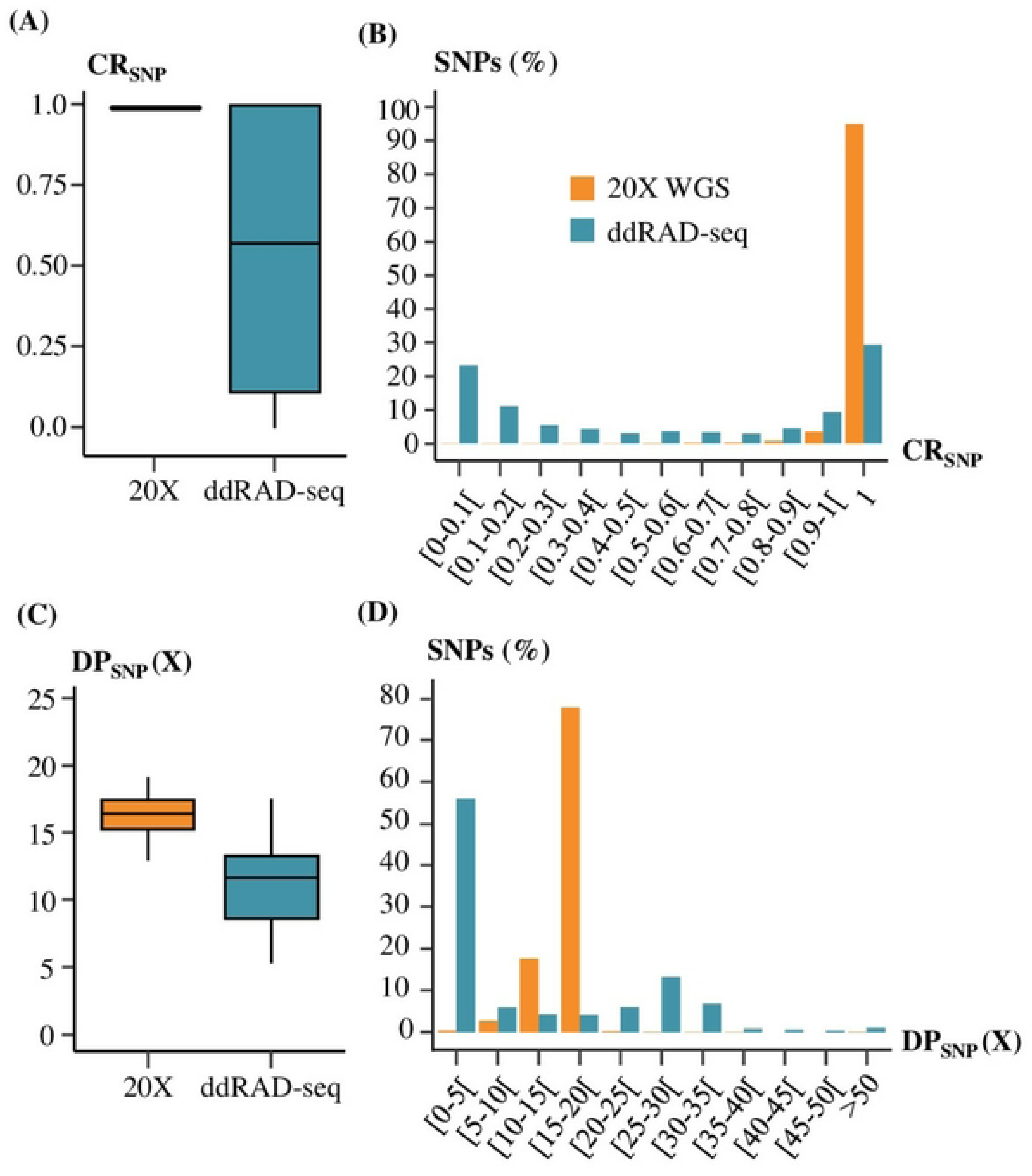
Comparison of variant calling results between 20X WGS (*in orange*) and ddRAD-seq (*in blue*). **(A) Distribution of CR_SNP_ values for 20X WGS and ddRAD-seq. (B) Percentage of SNPs per Call Rate SNPs (CR_SNP_) categories. (C) Distribution of DP_SNP_ values for 20X WGS and ddRAD-seq. (D) Percentage of SNPs per SNPs mean depth of sequencing (DP_SNP_) categories.**

Similarly to CR_SNP_, the average DP_SNP_ in ddRAD-seq was lower (11X) than the average DP_SNP_ observed in 20X WGS (16X) and even lower than the average DP_SNP_ expected (∼45X, Fig. 4C). The distribution of SNPs in DP_SNP_ categories showed two peaks of density: one at low DP_SNP_ values (0-5X) and one at higher values (25-30, Fig. 4D). Moreover, maximum DP_SNP_ values were 2,807X for 20X WGS and 547X for ddRAD-seq. But it was observed that 25% of the deepest ddRAD-seq DP_SNP_ values were superior to 24X X compare with 17X in 20X WGS (Fig. 4D). 56.0% of the ddRAD-seq SNPs were genotyped with less than 5X on average, against 0.5% in 20X WGS (Fig. 4D).

### Variant calling parameters at EFs scale

Theoretically in ddRAD-seq, 285,004 EFs between 200 and 500 bp should be generated and pair-end sequenced on an average of 150 bp. Therefore, we estimated that 85.5 Mb should be covered by ddRAD-seq. In laying hens, the genome size is 1.26 Gb (45) which means that we expect a mean DP_SNP_ close to 45X in ddRAD-seq (450 Gb per Novaseq 6000 flowcell with 120 individuals per flowcell).

For the SNPs inside and outside of the expected EFs, the CR_SNP_ and the DP_SNP_ were calculated. These two parameters were lower for the set of SNPs outside the theoretical EFs then for the SNPs inside the expected EFs (Fig. 5). Mean CR_SNP_ was 0.74 for SNPs inside the expected EFs and 0.25 for SNPs outside. The standard deviation of CR_SNP_ was 0.35 for ddRAD-seq SNPs inside and 0.27 for SNPs outside the expected EFs. These values appeared consistent with the peaks observed at a genomic scale (Fig 4B).

**Fig 5:**
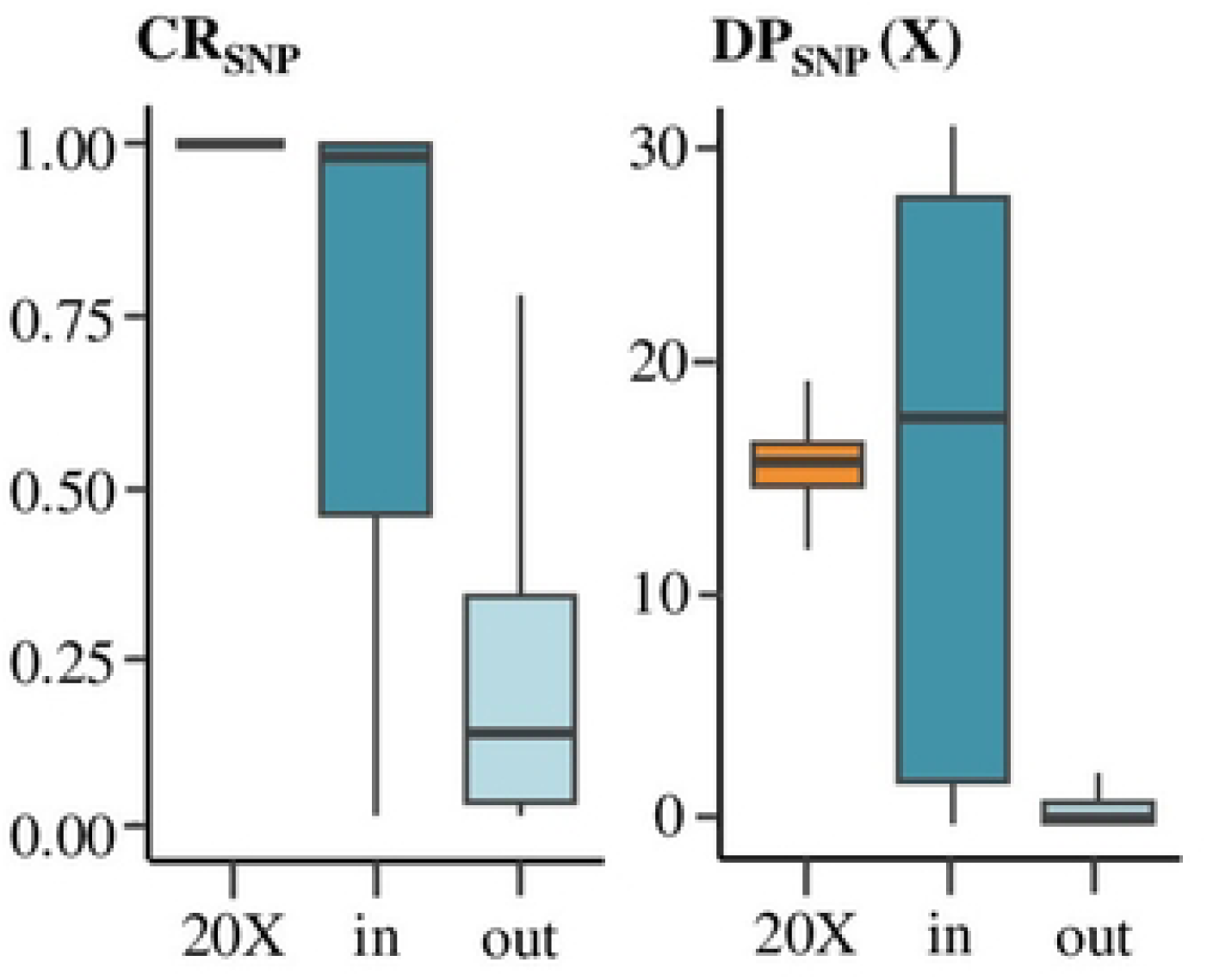
Distribution of CR_SNP_, DP_SNP_ between SNPs in (in blue) and out (light blue) of enzymatic fragments for ddRAD-seq compared to 20X WGS (in orange)

Mean DP_SNP_ values were respectively 17X and 1X for SNPs inside and SNPs outside the expected EFs (Fig 5). The standard deviation of DP_SNP_ for ddRAD-seq SNPs inside the expected EFs was 15X while it was 3X for SNPs outside. These DP_SNP_ values also corresponded to those observed for each peak at a genomic scale (Fig 4D).

Individuals mean depth of sequencing (DP_ind_) were 16X for 20X WGS and 11X for ddRAD-seq. Individual call rate (CR_ind_) were respectively 99.4% for 20X WGS and 55.4% for ddRAD-seq.

### ddRAD-seq Genotyping reliability

The concordance of ddRAD-seq genotypes with 20X WGS was assessed by comparing, when it was possible, ddRAD-seq genotypes to 20X WGS ones. Genotypes were said *comparable* when, for an individual, the SNP was genotyped by both ddRAD-seq and 20X WGS. Then, 9,398,316 genotypes were identified as comparable.

When comparable genotypes were identical between ddRAD-seq and 20X WGS, ddRAD-seq genotypes were said *concordant* with 20X WGS ones. Globally, 90.0% of the comparable genotypes from ddRAD-seq and 20X WGS were concordant. Most of these concordant genotypes (76.7%) were those of SNPs located inside the expected EFs. The other 13.3% of the concordant genotypes were those of SNPs located outside the expected EFs.

Discordant genotypes between ddRAD-seq and 20X WGS represented 10.0% of the total number of comparable genotypes. Among these discordant genotypes, 6.1% were genotypes from SNPs located inside the expected EFs. The other 3.8% of the discordant genotypes were from SNPs located outside the expected EFs.

The majority (7,784,818) of all comparable genotypes were those of SNPs located inside the EFs expected to be sequenced by the ddRAD-seq protocol (Table 2). Genotypes were more concordant between ddRAD-seq and 20X WGS for SNPs inside the expected EFs (92.6%) then for SNPs outside the expected EFs (77.7%).

**Table 2:** Effectives of comparable genotypes (GTs) according to the location of the associated SNPs, inside or outside the expected enzymatic fragments (EFs) theoretically generated by the ddRAD-seq protocol. The effectives of ddRAD-seq GTs matching (concordant) or not matching (discordant) with 20X WGS genotypes, when compared one by one for an individual, for a SNP.

|  | *Inside the expected EFs* |  | *Outside the expected EFs* |  |
| --- | --- | --- | --- | --- |
|  | *Concordant* | *Discordant* | *Concordant* | *Discordant* |
| *Number of comparable GTs* | 7,209,100 | 575,718 | 1,253,604 | 359,894 |
| *TOTAL* | 7,784,818 |  | 1,613,498 |  |
|  | 9,398,316 |  |  |  |

We also observed that the concordance between genotypes in ddRAD-seq and in 20X WGS, and the DP_GT_ of those genotypes were linked. Whether inside or outside expected EFs, concordant genotypes between ddRAD-seq and 20X WGS had greater DP_GT_ (15X) than discordant genotypes (7X, Fig. 6A).

**Fig 6.**
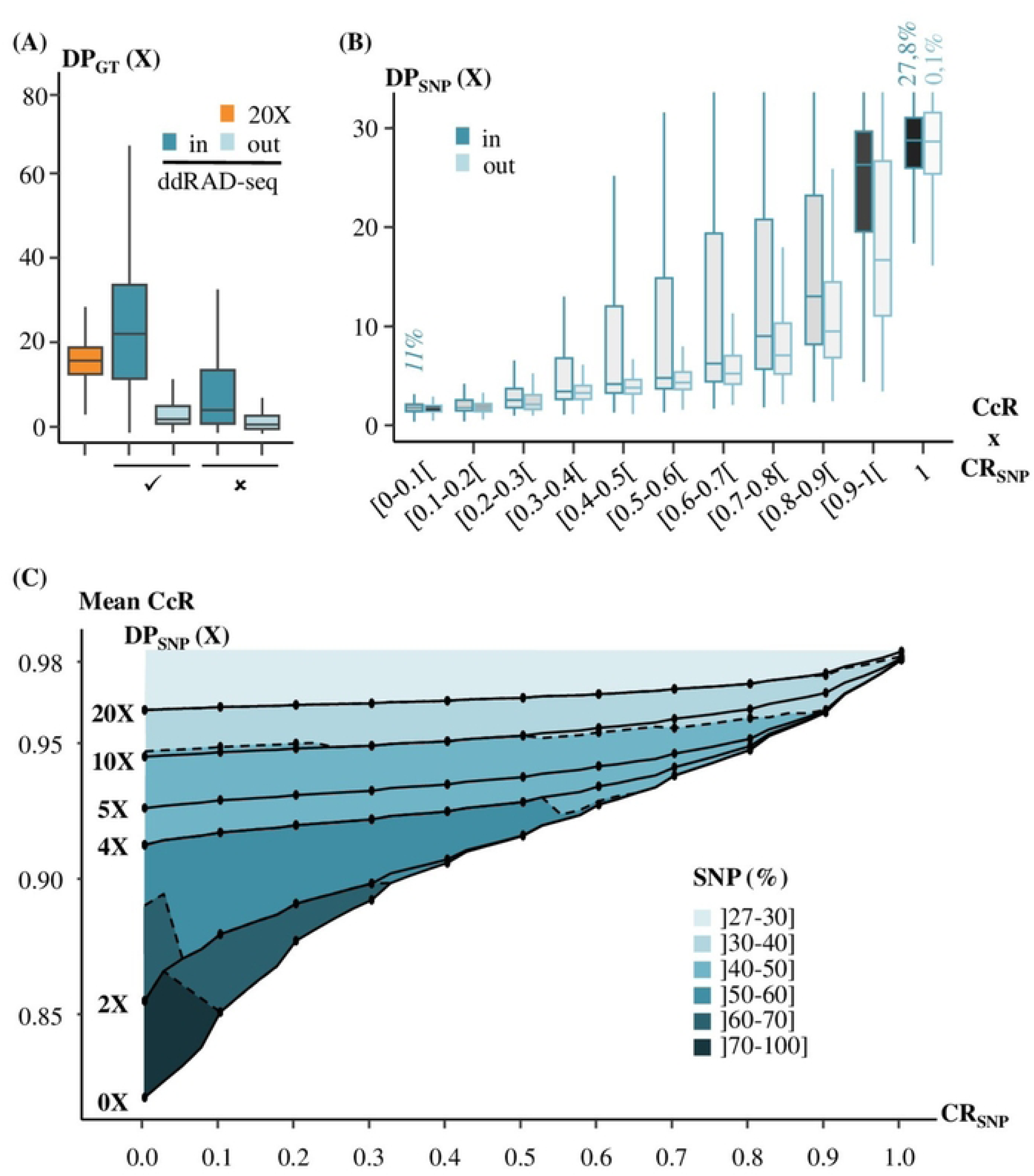
Genotyping quality of ddRAD-seq is correlated with the CR_SNP_ and the DP_SNP_. ***(A) Distribution of genotypes sequencing depth (DP_GT_) for 20X WGS, the reference, (in orange) and concordant (****✓**) or discordant (**✗**) genotypes in ddRAD-seq for SNPs in the enzymatic fragments theoretically generated by the ddRAD-seq protocol (in blue) or out the EFs theoretically generated by the ddRAD-seq protocol (in light blue). Percentages of each category of ddRAD-seq genotype on the whole amount of genotype was displayed (grey shades). (B) Distribution of SNPs sequencing depths (DP_SNP_) according to their category of genotype concordance rate (CcR) and SNP call rate (CR_SNP_). (C) Mean CcR of ddRAD-seq according to the CR_SNP_ and the mean sequencing depth threshold. The blue gradient represents the proportion of ddRAD-seq SNPs kept according to each filter combination***.

Then, for a SNP, the number of individuals with identical genotypes between ddRAD-seq and 20X WGS were quantified to calculate a concordance rates per SNP (CcR). The mean CcR for all the SNPs was 80.0%. The mean CcR was 87.3% for SNPs inside the expected EFs and 72.7% for SNPs outside the expected EFs. The correlation of CcR with the CR_SNP_ and the DP_SNP_ was investigated. High CcR values were associated with SNPs with high CR_SNP_ and DP_SNP_ values (Fig. 6B). Most of the SNPs with high CcR, CR_SNP_ and DP_SNP_ values were located inside the expected EFs. Inside the expected EFs, 27.8% of the SNPs were correctly genotyped for all individuals (CR=1 and CcR=1) with a mean DP_SNP_ of 29X. Outside the expected EFs, only 0.1% of the SNPs were genotyped with a DP_SNP_ of 26X (Fig 6B).

Finally, ddRAD-seq SNPs were filtered according to multiple combinations of DP_SNP_ and CR_SNP_ threshold. Then, the mean CcR for remaining SNPs and the percentage of ddRAD-seq SNPs kept after applying these filters were calculated (Fig 6C). Then, this multi-criteria filtering approach (DP_SNP_, CR_SNP_) makes it possible to assess the number of SNPs retained depending on the objectives of genotyping reliability. For example, by filtering ddRAD-seq SNPs according to a threshold of 5X for the DP_SNP_ and a CR_SNP_ of 0.8 in ddRAD-seq, approximately 40-50% of ddRAD-seq SNPs will be retained, and 95.1% of these retained SNP will have a concordant genotype with 20X WGS. With the application of QC filters on ddRAD-seq data (CR_SNP_ and DP_SNP_) with the highest thresholds possible, poor-quality SNPs can be eliminated by retaining a minimum of 27% genotyped SNPs for all individuals with 98% of reliable genotypes.

## Discussion

The aim of the study was to assess the variant calling and genotyping quality of ddRAD-seq in laying hens, as an alternative to low-density sequencing methods (LD chip or low-depth WGS). Due to the diverse nature of ddRAD-seq protocols (enzyme pairs, size filtration method, and bioinformatics processing), estimating the quality of our variant calling and genotyping data by comparing them to the literature was challenging. Therefore, the results from ddRAD-seq were compared to results from 20X WGS obtained for the same individuals. Prior research has demonstrated that comparing a test sequencing method to a reference method allows for estimating the quality of variant calling and genotyping (25). Additionally, for WGS, an effective coverage of 15X is considered as sufficient to achieve high-quality genotyping (2). With an observed average DP_SNP_ of 16X for the 20X WGS, it was deemed a reliable basis for comparison with ddRAD-seq and representative of reality.

Initially, the study revealed that ddRAD-seq exhibited high precision (93.7%). Literature reports precision rates for RAD-seq studies ranging around 90 to 94% (25). The SNPs were well-distributed across all chromosomes, including micro-chromosomes. This represented an advantage for ddRAD-seq compared to commercial chips that do not provide information on all micro-chromosomes. The genome-scale sensitivity of ddRAD-seq at 3.6% was comparable to similar methods found in the literature (0.5 to 5.5 % depending on enzyme pairs) (16).

However, at the scale of the theoretically expected enzymatic EFs by the protocol in our population, a significantly lower sensitivity of ddRAD-seq than expected was observed (32.4%). Half of the SNPs detected by ddRAD-seq were located outside the expected EFs, indicating that the sequenced EFs did not correspond to reality. This discrepancy is due to in silico estimations based on the known reference genome in the literature (6,46). To approach the actual digestion results and leveraging data from the 20X WGS, restriction sites created and destroyed by mutations were integrated from the outset. These mutations, documented as significant sources of variability in generated EFs among individuals, were described in the literature. It is also acknowledged that the quality of the reference genome significantly impacts in silico simulation of EFs (33). To mitigate this known bias, the latest version of the reference genome was used (47). Despite these precautions, a substantial difference between the expected REFs, based on protocol descriptions, and reality was observed.

Some SNPs were found in genomic regions covered by EFs smaller than 200 bp, despite the protocol’s filtration step. Literature describes that for ddRAD-seq, depending on the enzyme pair and size filtration method chosen, a significant difference in size distribution between the expected and sequenced EFs. This holds particularly true when using a 4-base cutter and a 6-base cutter as the enzyme pair, as in our case (46). Therefore, it is not surprising to find some SNPs in genomic regions covered by EFs smaller than 200 bp with ddRAD-seq.

We also noted a small proportion of SNPs (14.0%) located outside the anticipated EFs, within a 10-base proximity of the expected EFs. Drawing from existing literature, we hypothesized that these might have originated from degraded DNA fragments. Specifically, ddRAD-seq is highly sensitive to DNA quality among RAD-seq methods (8), known to significantly influence enzymatic digestion efficiency and susceptibility to UV light exposure (11). Some EFs may have been cleaved by a restriction site on one end and, despite the use of sticky end sites during adaptor ligation, improperly bound to these adaptors on the other end. Usually, after adaptor linking, EFs proceed through the rest of the protocol for amplification and sequencing. Based on previous observations in the literature, it’s plausible that despite the protocol’s trimming step, certain EFs smaller than twice the size of a read (∼300 bp) might have been sequenced in pair-end, potentially causing partial adaptor contamination (46).

Among the remaining SNPs that didn’t align with expected EFs or the previously described scenarios, some were genotyped by ddRAD-seq in genomic regions corresponding to EFs generated by the same restriction enzyme on both ends. This suggests that some EFs weren’t appropriately filtered to have two distinct adaptors at each end. According to the protocol, for an EF to be sequenced, it must be produced by the combination of Taq1 and Pst1. The majority of SNPs outside the EFs were situated in regions that corresponded to EFs cut by the same restriction site at both ends and were less than 200 bp long. We thus inferred that, within our ddRAD-seq protocol, the adaptor filtering step might not have been completely efficient. Several studies have also observed this phenomenon (48,49). These studies describe the presence of EFs generated exclusively by the first introduced RE in the protocol but not by the second one. In our case, we observed SNPs located in both Taq1-Taq1 and Pst1-Pst1 EFs with more Pst1-Pst1. Regarding the frequency of Taq1 and Pst1 RS on the reference genome, we hypothesized that the difference between the number of SNPs located in Taq1-Taq1 EFs or Pst1-Pst1 EFs was more due to the larger number of Pst1 RS. It is possible to assess the efficacy of the adaptor filtration step for EFs using methods like qPCR, yet very few studies do so (33,49), and they didn’t report the results. Similar to most studies employing ddRAD-seq, no quality control for this step was performed in our study. Therefore, it’s conceivable that the retention of a portion of EFs generated by a single restriction enzyme despite the filtration step is a generally acknowledged characteristic of ddRAD-seq.

Nevertheless, the random nature of previously cited bias leading events, among the pool of samples, should lead to average CR_SNP_ and DP_SNP_ values lower than those of the SNPs genotyped in the regions covered by the expected EFs. Globally, this would result in a loss of CR_SNP_ and DP_SNP_ at a genome-wide scale (11).

SNPs outside the expected EFs exhibited lower CR_SNP_ and DP_SNP_ compared to SNPs within the EFs, impacting the overall average values of CR_SNP_ and DP_SNP_ more negatively than anticipated. The sequencing depth intended for ddRAD-seq, originally allocated to specific regions, was spread across a larger area than expected. The theoretical calculation of wrongly assumed that only the theoretical EFs had been sequenced by the protocol. As a result, the average DP_SNP_ value decreased from the expected 45X to an observed 11X. Given that a minimum of 30X is recommended for ddRAD-seq, based on a reference genome, to ensure comprehensive genotyping in all individuals, the decrease in CR_SNP_ for SNPs called by ddRAD-seq was expected (18). As DP_SNP_ and sensitivity are correlated (1,50), this decline in DP_SNP_ is responsible for the low sensitivity observed at the expected EFs. Considering that low CR_SNP_ and DP_SNP_ values can impact genotyping reliability (51), it was hypothesized that genotyping SNPs outside the expected EFs might be less reliable than those within.

Upon comparison, the genotype concordance between ddRAD-seq and 20X WGS at 90% was highly satisfying. SNPs with high rates of erroneous genotypes (CcR) were indeed associated with lower CR_SNP_ and DP_SNP_ values than reliable genotypes.

Having genotype data for both ddRAD-seq and 20X WGS was a significant asset for our study, allowing for a detailed individual-level analysis of ddRAD-seq genotyping by pairwise genotype comparison. Quantifying genotyping errors enabled a thorough examination of the most common quality control filters’ impact on genotyping reliability. Many studies apply consensus threshold filters from the literature without quantifying their impact on genotyping reliability, which, depending on the applications, can significantly affect study conclusions (11,22).

Our study described the relationship between genotyping reliability and commonly used quality control filters for ddRAD-seq data (CR_SNP_ and DP_SNP_), while measuring their impact on the retained SNP quantity. Fig 6C allows for comparison among different quality control scenarios’ impact on genotyping reliability in terms of genotype reliability and SNP quantity. It offers the opportunity to establish decision rules on quality control thresholds tailored to each study’s needs.

Applying the most stringent quality control filters on CR_SNP_ and DP_SNP_. According to the literature, this marker count is largely sufficient for various applications in laying hens (35) such as genome-wide association studies (40), linkage disequilibrium calculation (41), CNV detection (42) or genomic selection (43). Hence, ddRAD-seq proves to be a reliable tool for laying hen genotyping, offering a superior number of SNPs with reliable genotypes, evenly distributed across the entire genome, making it a compelling alternative to LD chips and LD WGS.

## Materials and methods

### Animals

All animals consisted in a commercial pure line of laying hen of Rhode Island. This line was created and selected by *Novogen* (Plédran, France). The population studied was constituted of 50 roosters from the same generation, bred in individual cages.

### Whole genome sequencing

All 50 individuals were sequenced by the Genomics and Transcriptomics platform GeT-PlaGe (Toulouse, France) with the Illumina HiSeq2000 technology expecting a global coverage of 20X Firstly, 38 individuals were sequenced as part of UtOpIGe project. Secondly, 12 individuals from the project OptiSeq were sequenced. These individuals were chosen because they have been selected as breeders for further generation. Data were aligned to the GRCg7b chicken reference genome (47) with BurrowsWheeler Aligner V0.7.15 (52) with default parameters for paired-end alignment. SNP calling was performed with GATK V3.7 (53). Bi-allelic SNP have been extracted with the SelectVariant function and the “—restrictAllelesTo BIALLELIC” option. Remaining SNP have been filtered using the “VariantFiltration” option and hard filters for DNA-sequencing “FS > 60.0”, “QD < 2.0”, “MQ < 40.0”, “MQRankSum < -12.5”, “ReadPosRankSum < -8.0” and “SOR > 3.0”.

### ddRAD-sequencing

The same 50 individuals have been also sequenced with the ddRAD-seq technology as described in (10) by the **Montpellier GenomiX facility (MGX, France)**. Enzymatic digestion was performed using enzymes Taq1-v2 and Pst1-HF (New Englands Biolabs, 20assachusetts, USA), in agreement with the simulation results of Herry et al (2023). Only fragments ranging from 200 to 500 bp were selected as it is the appropriate length for sequencing fragments with **Illumina’s** sequencing systems **and more precisely the Novasesq 6000** (54). Mapping and variant calling were carried out in the same way as for the WGS sequences with the exception of the HaplotypeCaller module of GATK V3.7: -drf DuplicateRead argument, which was added to keep duplicated reads, as it is one of the principles of ddRAD-seq method.

### Identification of genomic regions theoretically covered by ddRAD-seq

Genomic regions defined by the ddRAD-seq protocol were estimated on the reference genome GRCg7b (47). First, all Taq1 and Pst1 RS were identified on the reference genome thanks to R package *Biostrings* (55). Then, RS created by mutations were identified using SNPs detected in 20X WGS as the list of possible mutations within our population. Then, EFs between 200 and 500 bp were generated and only the first and last 150 bp were kept. SNPs called in these regions by ddRAD-seq or 20X WGS were identified using bedtools v2.30.0.

SNPs called by ddRAD-seq outside of these regions were identified and their locations were studied. The location of theses SNPs regarding the position of RS on the genome was empirically observed using IGV 2.7.2. Hypothesis about the possible reasons of the identification of SNPs outside expected EFs were made based on these observations and comparisons with the literature. SNPs were then classified according to these hypotheses and counted using bedtools v2.30.0.

### SNP calling summary

For both ddRAD-seq and 20X WGS, the number and the repartition of genotyped SNPs along the chromosomes were estimated and compared using R V4.0.4. The correlation between the length of the chromosomes and the number of SNPs found on them was performed with the method of spearman for both methods using R V4.0.4. The average distance between two adjacent SNPs in bp was calculated on the whole genome and for each chromosome.

Then, for both ddRAD-seq and 20X WGS, three filtering parameters at the population scale were computed: (*i*) the ratio between the number of individuals with a non-missing genotype for a SNP and the total number of individuals, called the SNP call rate (CR_SNP_), (*ii*) the minimum allele frequency (MAF) in the population and (*iii*) the SNP sequencing depth (DP_SNP_) at the population scale. The genotype sequencing depth (DP_GT_) corresponds to the number of reads supporting a genotype. DP_SNP_ is the sum of each genotype sequencing depth divided by the total number of genotyped individuals for this SNP. CR_SNP_ calculations were performed using Plink V1.9 (56) and DP_SNP_ using VCFtools V0.1.16 (57) and R V4.0.4 (58).

### ddRAD-seq variant calling sensitivity and precision

The variant calling sensitivity of ddRAD-seq was calculated as the number of SNP commonly called by ddRAD-seq and 20X WGS divided by the total number of SNPs called by 20X WGS. The variant calling precision of ddRAD-seq was calculated as the number of SNP commonly called by 20X WGS and ddRAD-seq divided by the total number of SNPs called by ddRAD-seq. Sensitivity and precision were also calculated considering only regions covered by expected EFs.

### 20X WGS and ddRAD-seq genotype concordance

First, the SNPs that were called with both 20X WGS and ddRAD-seq approaches were kept. The number and the repartition of these common SNPs on the chromosomes were analyzed. Then, individually, each genotype was compared between 20X WGS and ddRAD-seq. If one or both methods didn’t allow to genotype the individual for a SNP, the genotypes were considered **incomparable,** and the SNP was excluded for this individual. Concerning the **comparable** genotypes, they were considered **concordant** when both alleles were the same between ddRAD-seq and 20X WGS. On the contrary, if one or two alleles were different for a genotype between ddRAD-seq and 20X WGS sequencing methods, they were considered **discordant**. For the concordant and discordant genotypes, DP_GT_ were compared between ddRAD-seq and 20X WGS. They were obtained using VCFtools V0.1.16. Moreover, the Concordance rate (CcR) was computed as the ratio between the number of concordant genotypes and the number of comparable genotypes for each SNP. The mean CcR was calculated on all the common SNPs. Finally, ddRAD-seq data were filtered according to multiple combinations of Mean DP_SNP_ and SNP CR_SNP_ threshold. The evolution of the mean CcR and the number of retained SNPs were studied under these conditions.

## Data availability

Data is deposited in the European Nucleotide Archive (ENA) with the accession number PRJEB58821 and PRJEB71464.

## Ethics statement

All blood samples were carried out as part of the commercial and selection activities of *Novogen*. These animals studied and the scientific investigations described herein are therefore not to be considered as experimental animals per se, as defined in European Union directive 2010/63 and subsequent national application texts. Consequently, we did not seek ethical review and approval of this study as regarding the use of experimental animals. All animals were reared in compliance with national regulations pertaining to livestock production and according to procedures approved by the French Veterinary Services.

## Author contributions

Conceived and designed the experiments: SA LL FL. Acquisition of data: EG. Analyzed the data: MD SA FL. Wrote the paper: MD FL SL SA FD.

---

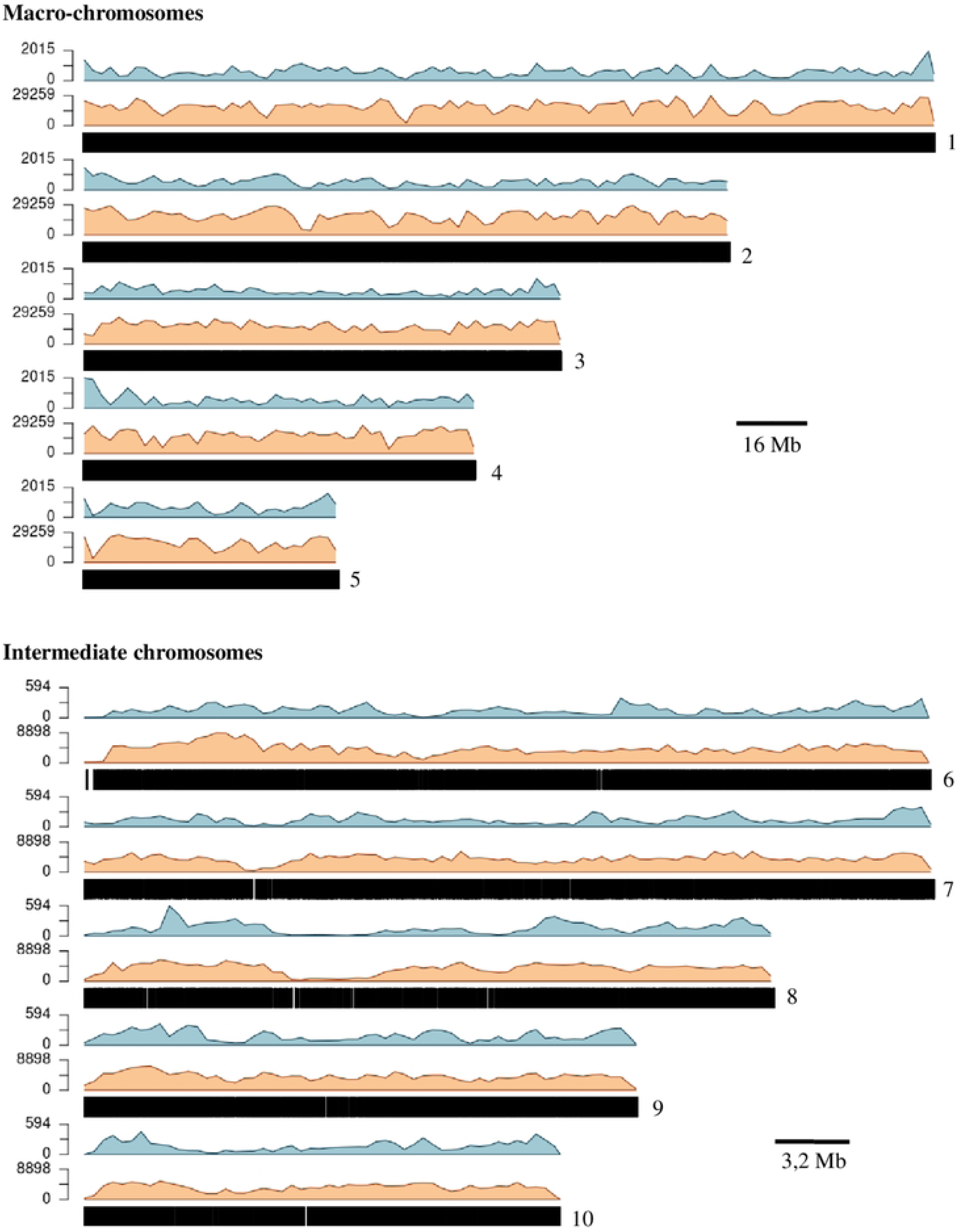

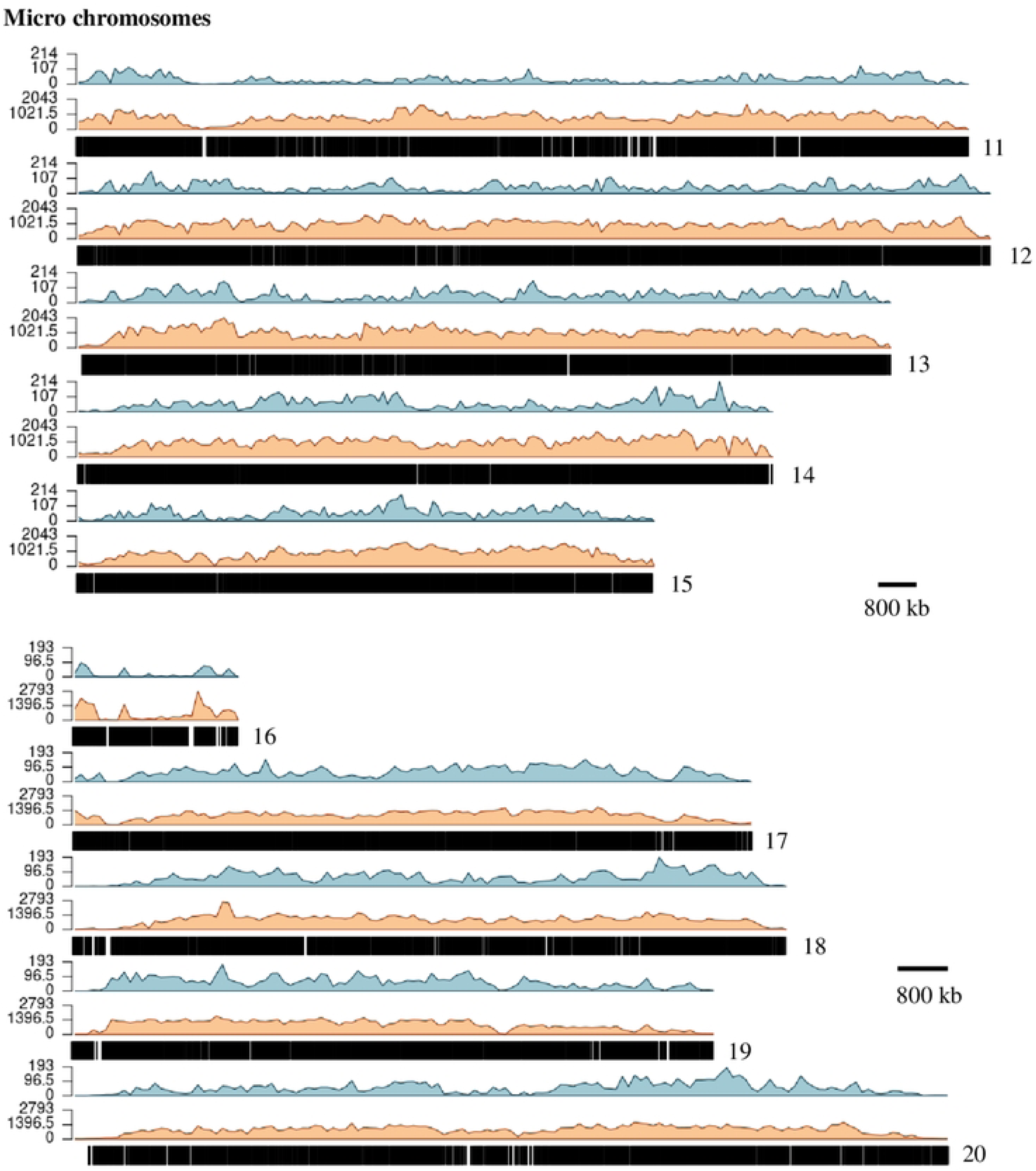

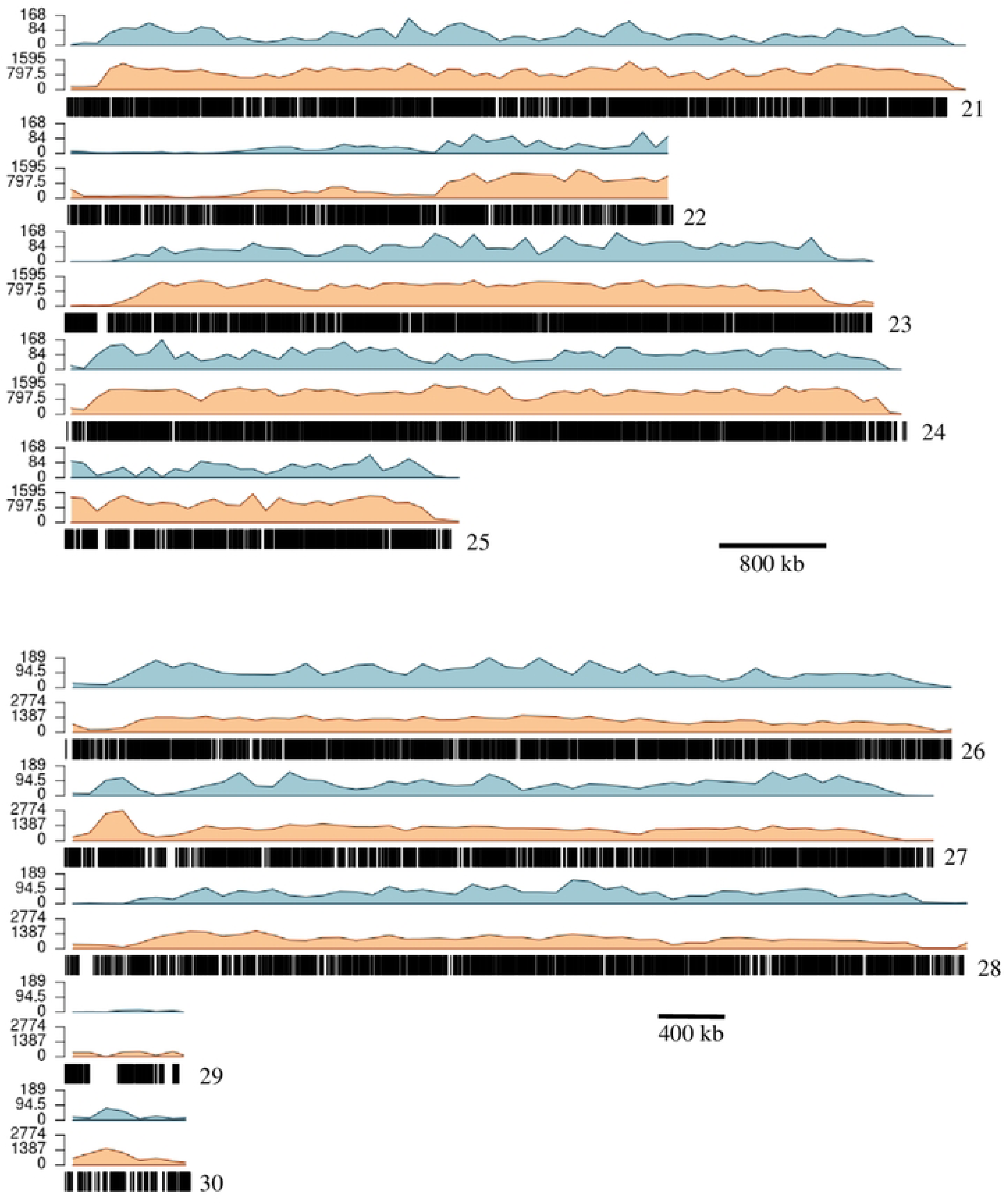

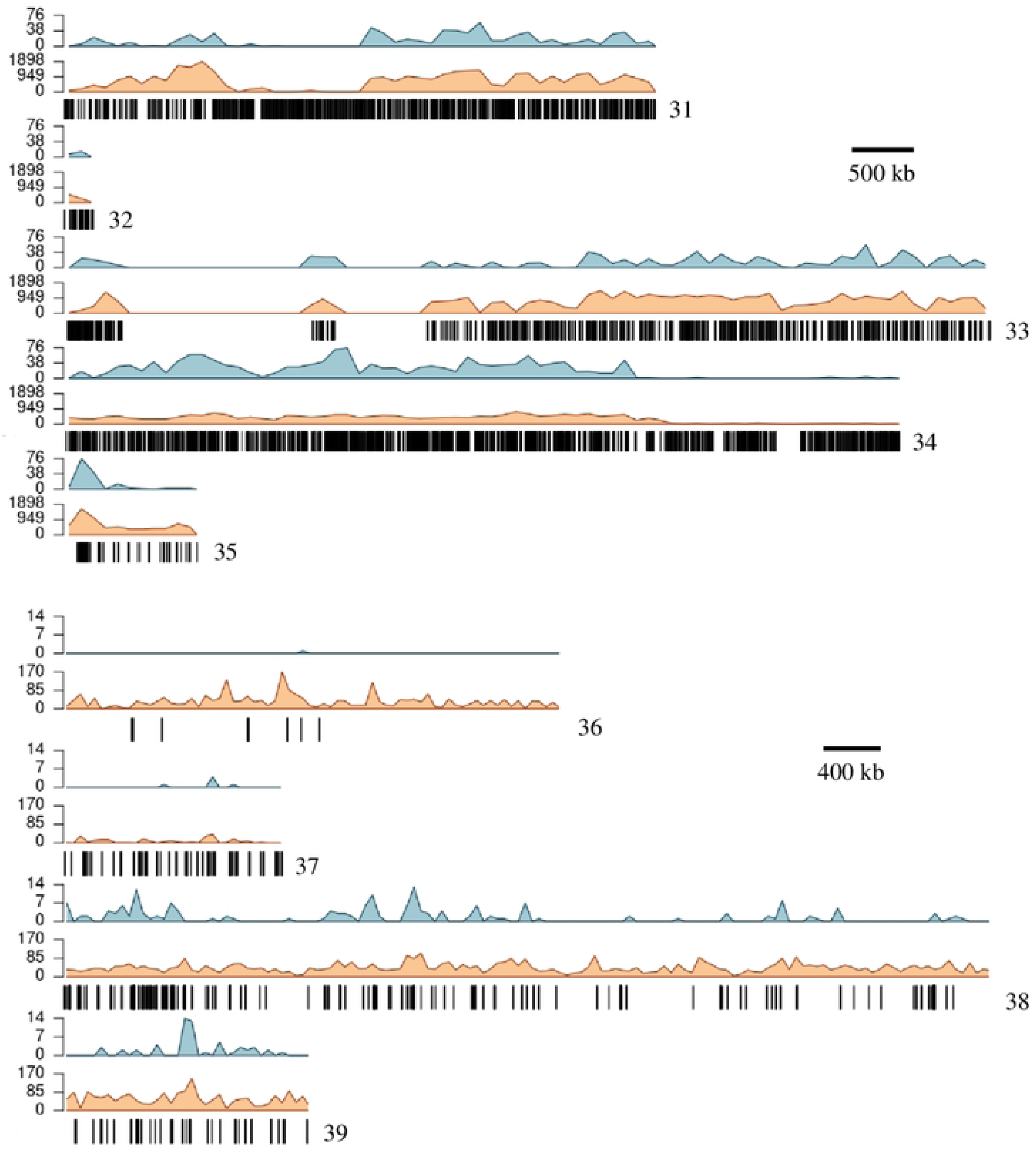

